# Using causality and correlation analysis to decipher microbial interactions in activated sludge

**DOI:** 10.1101/2021.09.26.461882

**Authors:** Weiwei Cai, Xiangyu Han, Hong Yao

## Abstract

Network theory is widely used to understand microbial interactions in activated sludge and numerous other artificial and natural environments. However, when using correlation-based methods, it is not possible to identify the directionality of interactions within microbiota. Based on the classic Granger test of sequencing-based time-series data, a new Microbial Causal Correlation Network (MCCN) was constructed with distributed ecological interaction on the directed, associated links. As a result of applying MCCN to a time series of activated sludge data, we found that the hub species OTU56, classified as belonging the genus *Nitrospira*, was responsible for completing nitrification in activated sludge, and mainly interacted with *Proteobacteria* and *Bacteroidetes* in the form of amensal and commensal relationships, respectively. Phylogenetic tree suggested a mutualistic relationship between Nitrospira and denitrifiers. *Zoogloea* displayed the highest *ncf* value within the classified OTUs of the MCCN, indicating that it could be a foundation for activated sludge through forming the characteristic cell aggregate matrices into which other organisms embed during floc formation. Overall, the introduction of causality analysis greatly expands the ability of a network to shed a light on understanding the interactions between members of a microbial community.

## INTRODUCTION

Ecological interactions, such as those involved in the exchange of resources or space, within microbial communities have been a topic of intense interest in microbial ecology, (Hibbing et al., 2010). The interactions of species are considered a driving force promoting ecological function of the microbial community, and due to its importance, the structure of communities have been described by species interaction networks for over a century (Berlow et al., 2009; Poisot et al., 2015). Although networks were initially was applied to the study of food webs, the concept has been expanded to microbial ecology to unravel ecological interactions (Ings et al., 2009; Kéfi et al., 2012). Therefore, microbial interactions within a community are more likely to be reflected by network theory, which can be established through a set of methodologies by mathematical correlation. Recently, network theory has been commonly used to explore the microbiomes of natural and artificial environments, such as soil (Barberan et al., 2012), sediments (Ji et al., 2016), bioreactors (Liang et al., 2018), and wastewater treatment plants (Global Water Microbiome Consortium et al., 2019).

In wastewater treatment plants, activated sludge has served as the core unit for wastewater treatment for over a century (Jenkins and Wanner, 2014). The highly diverse microorganisms in activated sludge thrive on organic compounds that are enriched in carbon(C), nitrogen (N), sulfur (S), phosphorus (P), and various trace elements, forming a complex web of ecological interactions based the competition for resources and space (Liébana et al., 2016; Xia et al., 2018). A series of graphical methods have been developed for constructing correlation or co-occurrence networks, to visualize and elucidate the complex microbial interactions of species in activated sludge, gut microbiome, natural environment (Weiss et al., 2016). Previous studies on co-occurrence or correlation networks have defined multiple relationships between species with a pairwise similarity matrix or sparse multiple regression analysis respectively (Faust and Raes, 2012). Generally, nodes and links in a network, respectively, represented species and interactions, yet, these interactions were only defined by positive or negative association, which limited further understanding of ecological interactions between species. As an intrinsic property of correlation analysis, previous networks were commonly undirected, demonstrating specific interactions among species, such as competition and symbiosis. Although a few studies have attempted directed networks, provided according to the time lag, to show a direction between nodes (Deng et al., 2016; Ju and Zhang, 2015), most studies rarely explore the possibility of causality analysis from time series data, which could enhance our understanding of ecological interactions.

Therefore, through a combination of correlation and causality analyses this study focused on constructing a directed network to discern the sophisticated interactions between members of an activated sludge microbiome. A previously published 259 day high-through sequencing data set was employed for correlation analysis and Granger test (Jiang et al., 2018). Coupling the correlation and causality analyses allowed construction of a microbial causal correlation network (MCCN), which demonstrated that the microbial interactions in activated sludge could be classified as mutualism, synergism, commensalism, neutralism, predation (parasitism), amensalism, and competition (antagonism). Hub-species OTU56 belonged to *Nitrospira* and showed more diverse interactions with *Proteobacteria* as compared to *Bacteroidetes*. Moreover, the *Zoogloea* were potentially the key genus that induced changes in many of the activated sludge bacteria due to its role in scaffold construction during sludge floc formation. The application of MCCN will provide information on the ecological interactions between different species in both natural and artificial ecosystems.

## RESULTS AND DISCUSSIONS

### Applicability of Granger causality

The assembly of the microbial community is commonly recognized as the result of deterministic and stochastic processes. The role of deterministic processes is considered to be limited in stable environments, therefore stochasticity could play an important role in gradually shifting community structure (Zhou and Ning, 2017). Due to the mutual influence of both processes, the abundance of a specific species is assumed to be the sum of a baseline and random variation. The variation of species over long periods of time should appear to be random. Within a steady state microbial community, the variation in abundance of specific species could be subjected to a joint distribution over time, as the present microbial community evolves from the previous state, while time should have limited influence on the variation of the microbial community. Although past observations are important to forecast future trends, these predictions do not completely depend on them. Therefore, there could be an autocorrelation process, which produces a time lag representing only finite past values that is applied to the forecasting. Deng et al. (2016) used the time lag to construct the correlation network with time-series data unravel microbial succession within an uranium bioremediation site (Deng et al., 2016). Additionally, David et al. (2014), when analyzing the effect of host lifestyle on human microbiota, relied on the autocorrelated process of time series (David et al., 2014), which demonstrated that OTUs variation complied with the time series model. We applied the data of 98 key OTUs obtained over the course of 259 days to fit into the ADF (augmented Dickey–Fuller test) test to verify whether microbial data is irrelevant to time or not. If the data is not stationary, which defined as the time series data is independent time, the difference between adjacent values will be applied to all data. The result of the stationary check is shown in Supplementary S2. All OTUs fulfilled the requirement of stationary after difference, 51 OTUs required difference treatment while the rest were stationary without difference treatment.

### Overall topological indexes of the causal network

The visualized causal network is shown in Fig 1. 98 OTUs were used for Microbial Granger Causal Network (MGCN) construction, which created 1865 links between the nodes at a significant threshold of p < 0.05. Granger causality is commonly not symmetric, network building had to be directed. The bidirectional links were defined as a feedback from the source to the target OTU, indicating that either node could improve the forecasting accuracy of the other. A unidirectional link indicates the source OTU significantly improved the forecasting accuracy of the target OTU but not vice versa. The outdegree and indegree directed links, defined by the direction of links in or out of the specific node, were counted separately. As shown in Table 2, the distribution of nodes degree tended to be normal rather than following power-law, regardless of whether indegree or outdegree, implying that the causal network was not scale-free (Deng et al., 2012).

**Figure 1.**
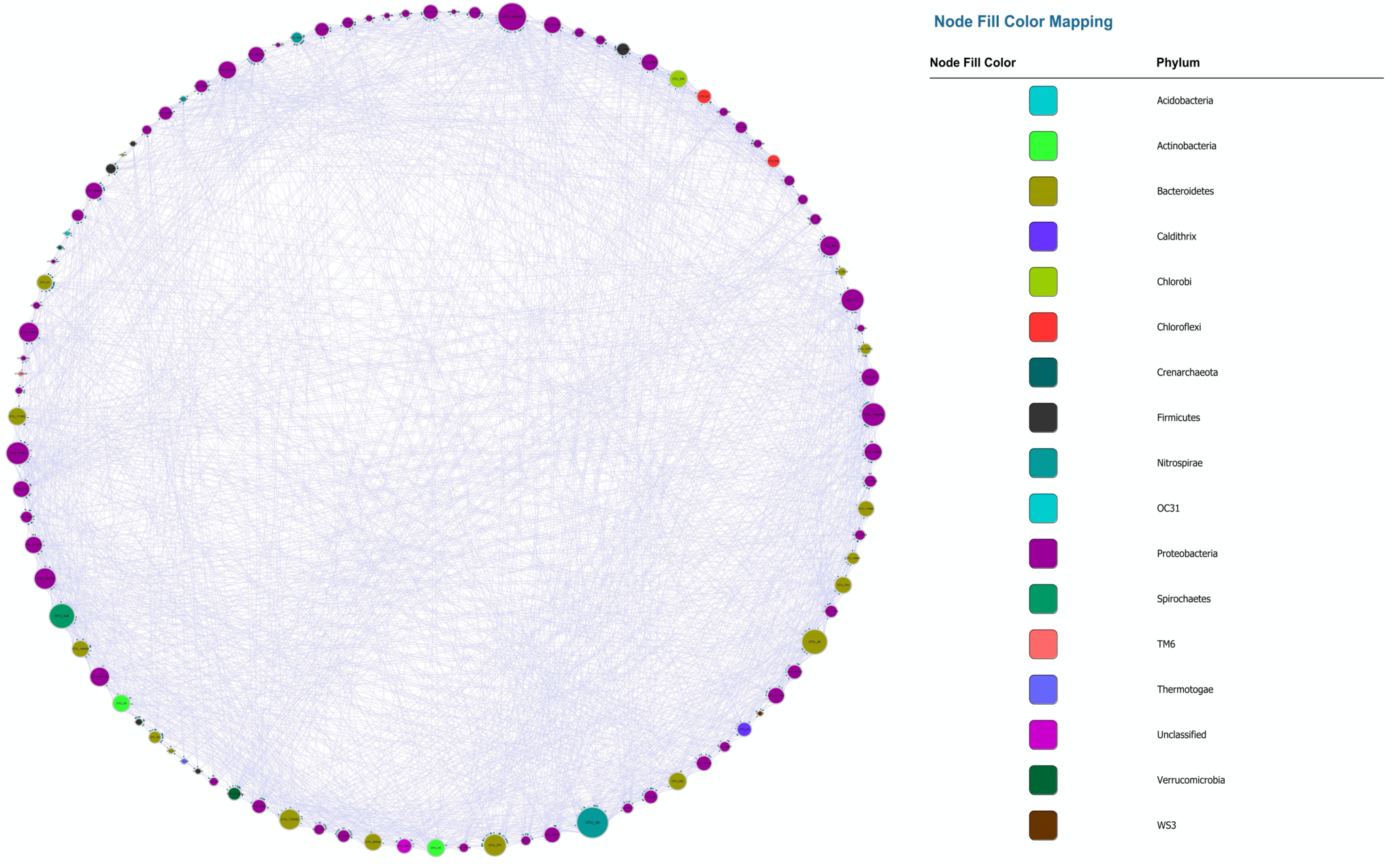
MGCN, each colour represents a separate phylum. The size of the node and node label is proportionate to the edge number of each node from 0 to 110. The arrows represent the direction of Granger causality.

**Table 1.**
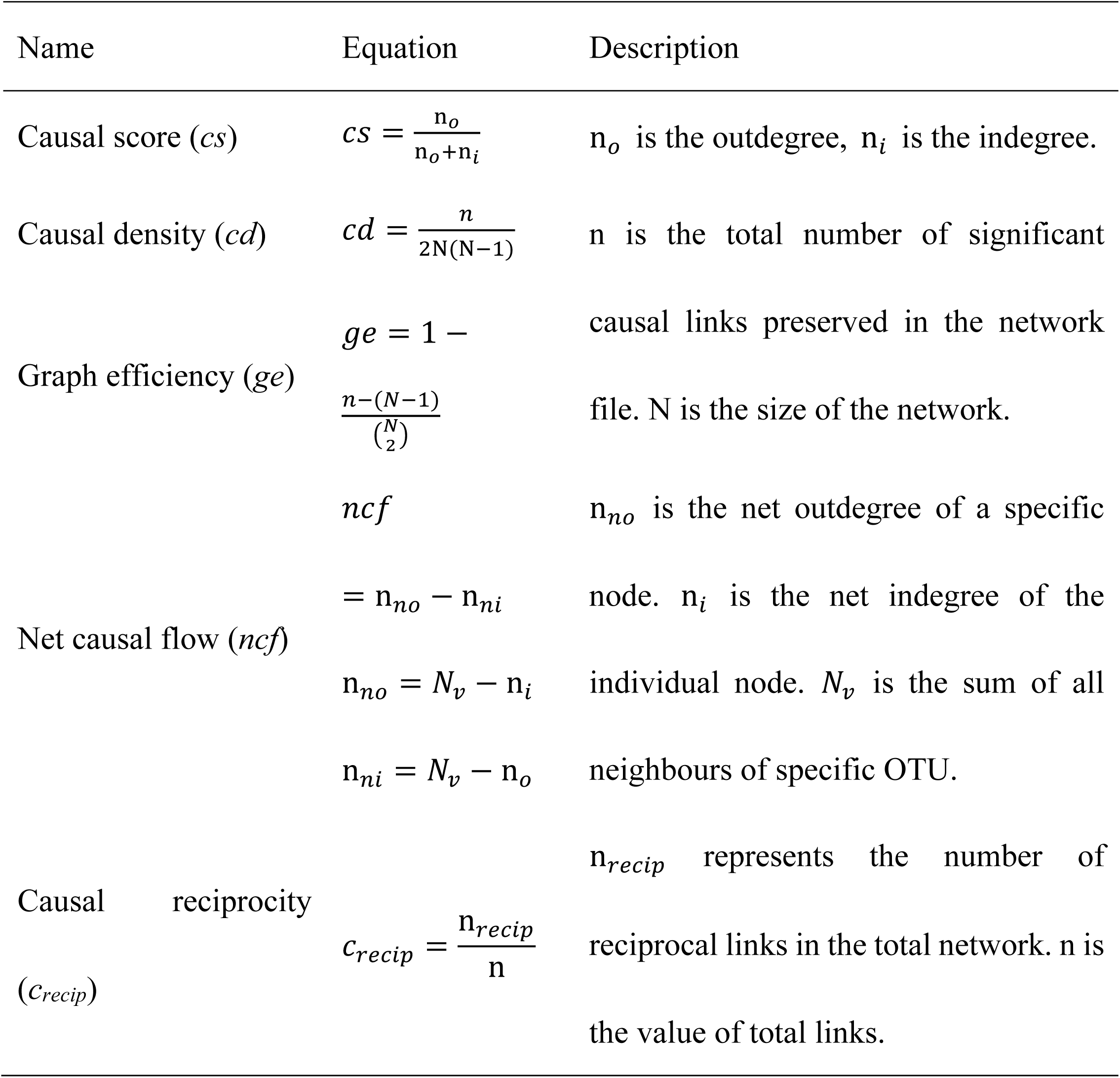
Network indexes

**Table 2.** Properties of different networks

|  | Granger |  | Random |  | Spearman |  | Random |  | MCCN | Random<br>MCCN |
| --- | --- | --- | --- | --- | --- | --- | --- | --- | --- | --- |
|  | MGCN | BoMGCN | MGCN | BoMGCN | MCN | BoMCN | MCN | BoMCN |  |  |
| network size | 98 | 81 | 98 | 81 | 98 | 98 | 98 | 97 | 73 | 73 |
| network density |  |  |  |  | 0.811 | 0.568 | 0.811 | 0.568 |  |  |
| links | 1856 | 730 | 1856 | 730 | 3856 | 2644 | 3856 | 2644 | 441 | 441 |
| power law (in) | 0.041 | 0.468 | 0 | 0.284 | 0.211 | 0.101 (in total) | 0.292 | 0.02 | 0.552 | 0.016 |
| power law (out) | 0.03 | 0.064 | 0.043 | 0 |  |  |  |  | 0.595 | 0.111 |
| average clustering coefficient | 0.449 | 0.373 | 0.196 | 0.119 | 0.868 | 0.753 | 0.812 | 0.57 | 0.352 | 0.084 |
| network diameter | 6 | 8 | 3 | 4 | 2 | 4 | 2 | 2 | 7 | 5 |
| network radius | 3 | 1 | 2 | 3 | 2 | 2 | 2 | 2 | 1 | 3 |
| average shortest paths | 2.149 | 2.647 | 1.823 | 2.21 | 1.189 | 1.463 | 1.189 | 1.432 | 2.866 | 2.571 |
| average number of neighbors | 27.082 | 14.716 | 34.143 | 17.21 | 78.694 | 54.515 | 78.694 | 54.515 | 9.753 | 11.507 |
| *cd* | 0.098 | 0.056 |  |  |  |  |  |  | 0.042 |  |
| *ge* | 0.815 | 0.9 | NA |  | NA |  | NA |  | 0.93 | NA |
| average *cs* | 0.491 | 0.504 |  |  |  |  |  |  | 0.509 |  |
| average *C<sub>recip</sub>* | 0.238 | 0.148 |  |  |  |  |  |  | 0.157 |  |

NA represents there is no data for this property.

The average clustering coefficient, which reflected the clustering degree of the overall network, was defined as the average of clustering coefficient over all nodes. The clustering coefficient of MGCN (0.449) was higher than previously described undirected networks, including grassland soils (0.1∼0.22), lake sediment (0.09), and groundwater condition (0.17-0.29), and was comparable with the value of 0.466 observed in a previous activated sludge study (Ju and Zhang, 2015). Watts and Strogatz (1998) introduced the random rewiring procedure to interpolate regular and random networks, in which the regular lattice is highly clustered while the random network is poorly clustered (Watts and Strogatz, 1998). Therefore, the higher relative clustering exhibited by the causality indicated that the network was defined rather than random. The average shortest average path was 2.149, which was smaller than within the undirected network. Hence, we derived a relatively clustered network connected by shorter paths, demonstrating that neighbouring nodes were closely connected. To confirm the small-world property, randomized networks with the same nodes and degrees as the original network were constructed. The average clustering coefficient and shortest paths were ∼0.196 and ∼1.823 respectively, whereas the ratio of Granger network to the random network of clustering coefficient and shortest path can be determined (Liao et al., 2011). As the ratio was equal to ∼1.943, this indicated the network possessed small-world properties.

### Indexes of nodes

According to the definition of *cs*, its magnitude represents the ability of a specific OTU to cause the variation among its neighbours. A value of 1 indicates that an OTU can affect its neighbours without being affected by them, while zero indicates the opposite. The *c*_*recip*_ reflected counts of reciprocating links, which exhibited feedback behaviour of each OTU, thereby higher values indicated that an OTU is likely to interact with others. Therefore, as shown in Fig. 2, as *c*_*recip*_ increases, the *cs* will tended to approach 0.5, displaying an equilibrium of indegree and outdegree links. All nodes displayed a *c*_*recip*_ value of less than 0.5, suggesting bidirectional links were not dominant in the relationship of all nodes. However, it was interesting that more interactions could be positively related to the equilibrium trend of *cs*. The *cs* and *c*_*recip*_ were both relatively quantified as the proportion excluded the magnitude of degrees, node size in Fig. 2 is proportional to degree of connection with neighbouring nodes. The average number of neighbours for a node was ∼27.08. The majority of nodes with a large number of neighbours had higher *c*_*recip*_ and neutral position of *cs*. Nodes with lower *c*_*recip*_ and lower *cs* indicated that more links were indegree, with the reverse, higher *c*_*recip*_ and *cs* indicating more links were outdegree. Integration of the relative proportion and neighbour number that was considered as an absolute quantity was beneficial for inferring the central output nodes in the network, which should possess lower *c*_*recip*_, higher *cs*, within fairly large size of neighbours. The average of *cs* was ∼0.491, showing that the number of outdegree and indegree links were nearly identical. The average of *c*_*recip*_ was 0.24, implying mutual cause is not predominant due to the lower proportion in total links. Additionally, *ncf*, the difference between net outdegree and net indegree, of nodes ranged from -20 to 21, as shown in fig. S3. The average of individual outdegree was 8.14, the average net indegree was the same. Moreover, the number of OTUs with positive *ncf* were greater than that of negative *ncf*, indicating more than 50% of the relationships in the system displayed Granger causality in the activated sludge system.

**Figure 2.**
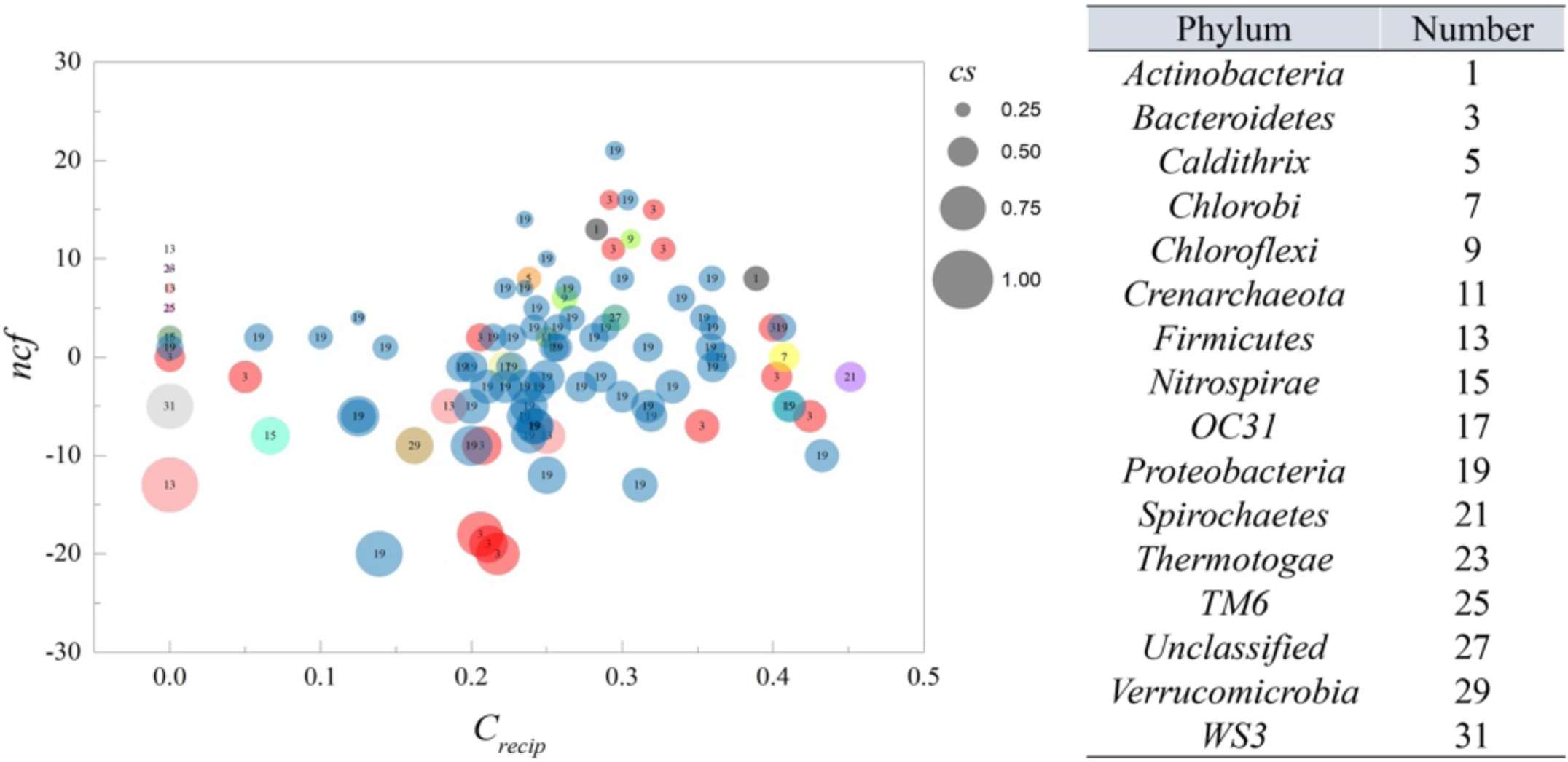
*ncf* and *c*_*recip*_ from MGCN. Each circle represents an individual node from MGCN with size representing the *cs* value. The number within the circle corresponds to the classification of OTU at the phylum level.

### Bonferroni-correction

The Bonferroni-corrected MGCN (BoMGCN) was produced from the original significant network (Fig. 3). The corrected network was sparser in comparison with the causal network, containing only 81 nodes and 730 links, and a lower clustering coefficient (0.373). The reduced network was highly conservative as the Bonferroni-correction excludes all potential type I error (false link was accepted) and displayed a slightly improved stability as revealed by the R square of power-law. The value of outdegree R square was 0.064, close to zero, yet the value for indegrees was 0.468, indicating a significant increase. Although the values were too small to be wholly fitted into power law, they indicated that some nodes in the BoMGCN had a greater or lesser effect on other nodes. An improvement of scale-free property was also observed, as well as an increase in the small world index, as represented by an increase in the ratio of σ (∼2.617), caused by a decrease in clustering coefficient and increase in the average shortest path, showing BOGCN is more likely to fall in the rules of a small world. Within the random network derived from BOGCN there was a clear decrease in clustering coefficient (0.119). Additionally, the properties of total nodes were slightly distinguished from the original MGCN network as a clear decline of *cd* value. The average *cs* increased from 0.491 to 0.504. Overall, the BoMGCN reduced the size of the network while keeping its basic properties. According to the classification of OTUs in BoMGCN, *Proteobacteria* was the predominant nodes, the hub species was *Nitrospira*, indicating the nitrogen-associated species has a broader social connection with other microbes.

**Figure 3.**
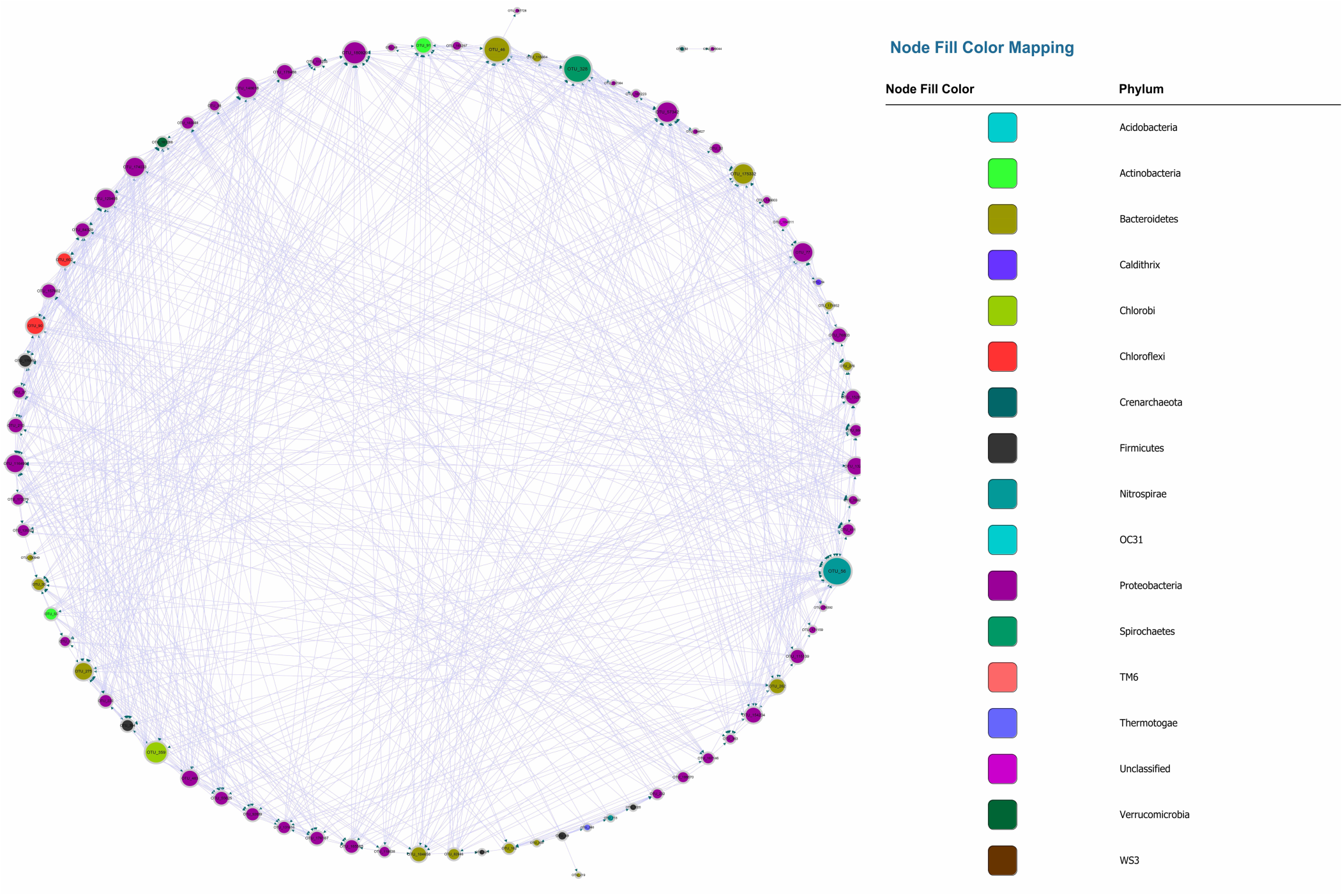
BoMGCN. Each colour represents a separate phylum. The size of the node and node label are proportionate to the edge number of each node from 0 to 110. The arrows represent the direction of Granger causality.

### Correlation-network supplemented to causal network

The MGCN showed the casual effect within the microbial community, however, information about positive or negative correlations between nodes was missing, therefore a Bonferroni-corrected microbial correlation network (BoMCN) based on Spearman’s correlation (shown in Fig. S4) was applied to supplement the MGCN, constructing a Microbial Causal Correlation Network (MCCN). The multiple relationships between two OTUs could be revealed more explicitly according to this combination of causality and correlation. Previously, correlation analysis was generally used to discern the negative and positive relations within a microbial network, indicating the ecological interactions between members of the community (Faust and Raes, 2012). As shown in Figure 4a, a combination of correlation and Granger causality could construct a new relationship, which shows the directional connection among nodes including the positive or negative effect they have on each other. As shown in Figure 4b, the MCCN was composed of 73 nodes and 441 links. Although the causality is at a higher level compared with correlation, i.e., all nodes with causal links should show strong mutual interaction, the missing nodes and links could be ascribed to the Granger causality, that is not a real causal relationship, due to the limitations of the method. Technically, the Granger test has been widely used for predicting the causal effect, in the context of the current study, the Granger test was utilized to forecast the relations between OTUs, which may lead to a better understanding of microbial behaviours and relationships within a community. The network can be used as an essential tool to predict microbial interaction when the real community could be too complex for accurate study, as additional efforts would be required to verify interactions between species.

**Figure 4.**
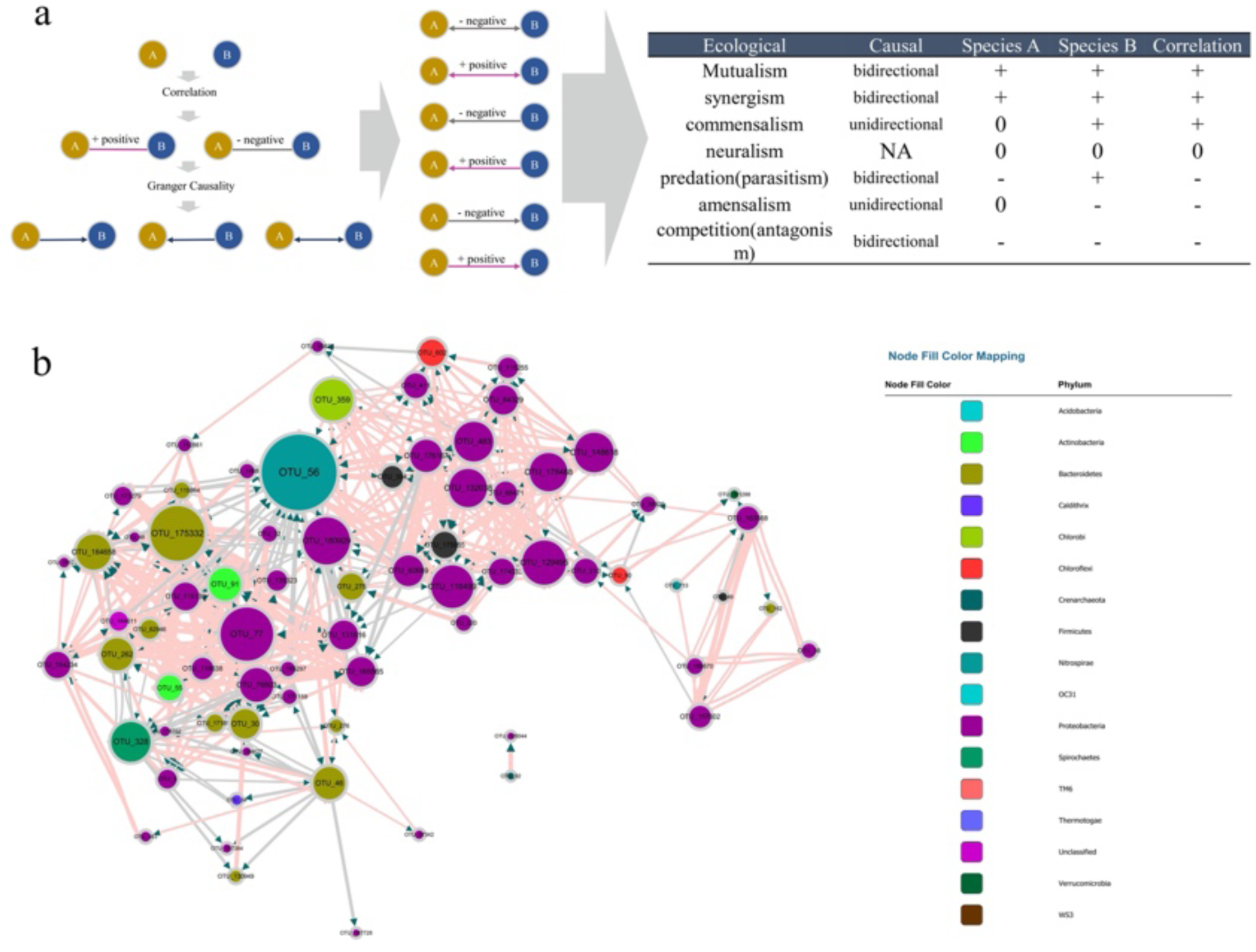
(A) Principle of the MCCN inference from the combination of causality and correlation, the detail of correspondence from MCCN link to ecological interaction on the right table. (B) MCCN, each colour represents an individual phylum. The size of the node and node label are linearly proportionate to the edge number of each node from 0 to 50. The arrows represent the direction of Granger causality. Pink and grey link colours represent positive and negative associations, respectively. The size of the link is proportionate to the correlation absolute value from 0 to 1.

The combination of correlation and Granger test allows observation of more specific interactions between two species, therefore a MCCN network could be applied to predict ecological relationships for community analysis. As shown in Figure 4a, there are seven patterns of species interactions, including mutualism, synergism, commensalism, neutralism, predation (parasitism), amensalism, and competition (antagonism) (Pepper et al., 2015). According to the results of MCCN, both mutualism and synergism should be a bidirectional edge with a positive effect on both species, as each species would derive benefits from the other, such that it would be difficult to distinguish them apart. Commensalism can be reflected by a unidirectional link with positive effect as species A can obtain a metabolite produced by species B. Although species B would be irrelevant to the growth of species A, as there is no feedback from A to B, the sequencing data of two species would be positively correlated as more species B would secret more metabolites for species A. Oppositely, a unidirectional connection with negative effect is classified as amensalism due to the general release of inhibitors from species A to species B. Here the quantity of species A will be relevant to the production of inhibitors, such as antibiotics, which can reduce the number of species B, thereby the contrary growth of species A and B will lead to a negative correlation. Although the predation (parasitism) can be implied by the negative bidirectional edge, the sequencing data used in this study contained only information from the 16S rRNA gene of bacteria, with no information about protozoa or phages, resulting in the exclusion of predation (parasitism) from the MCCN of the microbial community (Deng et al., 2016). Finally, a negative bidirectional link could also indicate competition between species. The MCCN is a powerful tool to recognize multiple interactions of microbes by specifying the endogeneity of correlation, which has been widely used as a statistic proof of microbial interaction within a network (Weiss et al., 2016).

### Core species in MCCN

The nodes with amounts of links would be considered as “hubs” in the MCCN. OTU56 was the hub species with the greatest number of indegrees (31) and second highest number of outdegrees (16). It was classified as belonging to the genus *Nitrospira*, a globally distributed group of nitrite oxidizers, which are capable of completing nitrification from ammonia to nitrate by one step (van Kessel et al., 2015). As shown in Figures 5, S5, and S6, OTU56 closely interacted with 24 OTUs from the phylum *Proteobacteria*, 8 OTUs from the phylum Bacteroidetes, and the 6 remaining OTUs interacted with 5 additional phyla. 21 OTUs displayed negative interactions with *Nitrospira*, 14 should be amensalism and 7 were competition relationship. Interestingly, all competition interactions originated from *Proteobacteria* to *Nitrospira*, showing a number of *Proteobacteria* may depress the growth of *Nitrospira*. This could be ascribed to the fact that most bacteria related to nitrogen cycle were *Proteobacteria* (Costa et al., 2006). Additionally, OTU56 unidirectionally interacted with OTUs from *Bacteroidetes*, for which there were only two types of interactions, commensalism and amensalism, with 3 and 5 links, respectively. According to a global diversity and biogeography study of over 300 wastewater treatment plants, only 28 out of 61448 OTUs, accounting for 12.4% of the 16S rRNA gene sequences, were defined as core OTUs, and these mainly consisted of *Proteobacteria, Bacteroidetes*, and *Nitrospira* in activated sludge (Global Water Microbiome Consortium et al., 2019). Therefore, the results of MCCN in this study are consistent, as *Proteobacteria* and *Bacteroidetes* actively interacted with the core species of *Nitrospira*, a group which plays a crucial role of nitrification in activated sludge. At the genus level, the majority of species that interacted with OTU56 were unclassified, and of those that could be identified, *Azospira*, which possesses denitrification activity, exhibited a mutualistic relationship with *Nitrospira*, as well as with OTU176167 and OTU92689, which were most closely related to the genus *Dechloromonas*, members of which are capable of reducing nitrate or chloride. The above mutualistic relationships could be achieved in nitrogen cycling processes, with denitrification removing nitrate as a product inhibitor to *Nitrospira*, meanwhile, *Nitrospira* could supply nitrate as a substrate for denitrifies.

**Figure 5.**
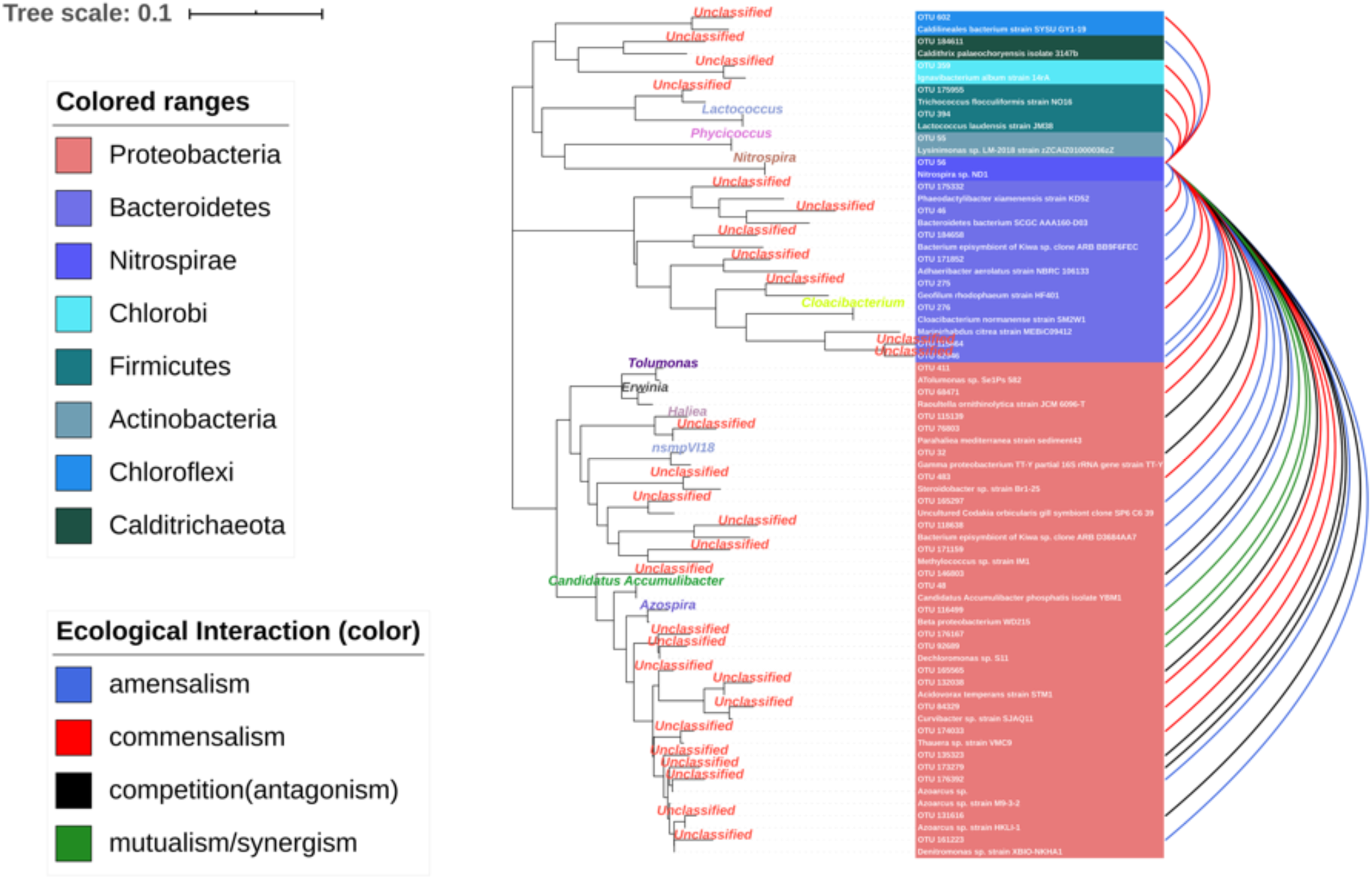
Ecological interaction of OTU56 with others at the OTU level. The colour represents the type of interaction. The phylogenetic tree shows the closest species according to the results of NCBI blast.

OTU180929, which had the most outdegree links (17), was classified as *Sinobacteraceae* at the family level. Members of this family are known to play a role in the degradation of aliphatic, aromatic hydrocarbon compounds and small organic acids (Gutierrez et al., 2013; Zhang et al., 2018). The number of net outdegrees and net indegree indicated the trending of nodes to cause a change of others or be affected by others. OTU180929, belonging to the genus *Zoogloea*, possessed 13 net outdegree and 13 net indegree links separately. *Zoogloea* has previously been demonstrated to be a bacterial genus important in the process of floc formation (Shao et al., 2009), and in this study is represented by OTU180929 and OTU178488. As shown in Figure S7, the *ncf* of *Zoogloea* was the highest value within the sum of classified OTUs, indicating that *Zoogloea* could enhance the growth of most species, i.e., it could be the foundation for the formation of activated sludge. However, the *ncf* of unclassified OTUs was still higher, reaching 18. The culture-depedent methods build the basics of microbiology research, which investigate the role of specific species (mostly are filamentous) in sludge flocculation and foaming (Nielsen et al., 2009). The unclassified nodes in MCCN showed there is still a massive microbial dark matter in activated sludge wait to be cultured. The network approach has been used to elucidate and prioritize the microbial dark matter in microbial community (Zamkovaya et al., 2021). Although activated sludge has been a widely employed strategy in wastewater treatment plants for over 100 years (Nielsen and McMahon, 2014), its microbiome still contains many mysteries, and is abundant with unknown species that are only gradually being elucidated by recent progress in culture-dependent and independent technologies.

In conclusion, the coupling of correlation and causality was crucial to understand ecological interactions within the microbial community. The Microbial Causal Correlation Network (MCCN) showed a sophisticated causal network in activated sludge and identified the fundamental species, with highest *ncf* value, as *Zoogloea*. The Microbial Causal Correlation Network (MCCN) and phylogenetic analysis together pointed out the core-species of *Nitrospira* (OTU56) could have mutualistic interactions with denitrifiers in activated sludge. However, most species that interacted with OTU56 were still unclassified, implying a greater sequencing depth would be the key to improve the understanding of activated sludge.

## MATERIALS AND METHODS

### Sequencing data derivation

The sequencing data were acquired from NCBI (accession number: PRJNA324303), which has been published previously (Jiang et al., 2018). The time-series data set included sequencing data for 259 days taken from a long-term operational wastewater treatment plant. The primers were F515 and R806, which covered mostly bacteria and archaea. The achieved fastq files were combined and processed online using a galaxy platform (Feng et al., 2017). OTUs were created with 97% cut-off through Uparse clustering method. RDP classifier assigned one representative sequence from each OTU to bacteria or archaeal taxonomy according to the 16S rRNA Greengene Database. The final OTUs table was prepared for the subsequent process. The phylogenetic tree was created with Mega software with N-J method, the visualization was completed online (https://itol.embl.de) (Letunic and Bork, 2019).

### Stationarity

Stationarity is an important concept to time series analysis and is a precondition to Granger Causality. The properties of stationarity were defined by the three main factors in terms of mean, variance, and covariance. The stationarity indicates that there was no change of trend in the time data, and it is known as a changeless process of the joint distribution within a specific displacement. The stationary implies that the expectation value of OTUs will fluctuate around the mean value of their neighbourhood rather than depend on time. This allows an estimation of the significant interval for the variation. Therefore, the stationarity analysis should be performed before analyzing time series data. It can be tested by detecting the presence or absence of unit root. The ADF-test was employed to verify if the time series data conformed to the stationary. If the original data is not subjected to stationarity, we used the difference, one minus another one to calculate the difference, to obtain the stationarity data. All abundance data of OTUs were filtered with the stationary test, while data that failed to go through ADF test after two rounds of using difference would be summed in the separate file as nonstationary data. Although the abundances of OTUs may vary on a large scale, even seemingly without a mean value, the difference would be stationary in most situations. The operating reactors could be affected by many factors, which would shift the microbial community via stimulation the metabolism of specific species.

### Granger Causality

The Granger causality test is a statistical hypothesis test that determined the role of one time series in forecasting another one (Granger, 1969). Herein, the Granger causality is limited within interpreting the interaction of two OTUs which were subjected to the ARMA (Autoregressive–moving-average model) model. To *i*th OTU, the ARMA model is shown as the equation:

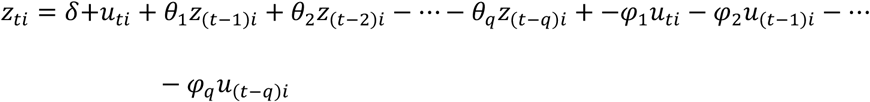

We simplified the equation for OTUs to the following format.

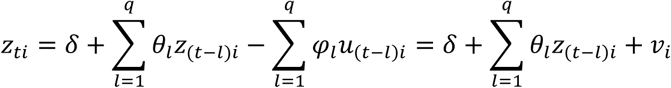

*v*_*i*_ is the random variation (white-noise series). Thus, we assumed the model for *i*th OTU is X, the model for *j*th OTU is Y. Both equations are as follows:

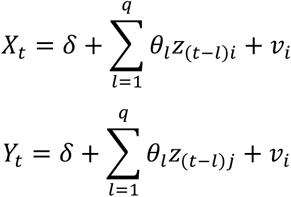

To know the interaction of X and Y, we assumed X and Y are interplays in their respective model predictions. The new models are derived as:

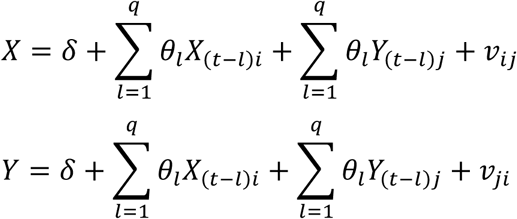

If the time series data followed the above equation, meaning the past values of X will contribute to predict current Y, and vice versa. However, real data could be applied to the following equation:

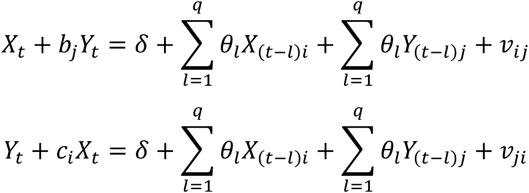

If *b*_*j*_ and *c*_*i*_ are not equal to 0 at the same time, this will be a model with instantaneous causality. In other words, the *v*_*ij*_ and *v*_*ij*_ would be the key to determine the Granger causality, if the variation could be decreased when applying the *b*_*j*_ ≠ 0, representing the *j*th OTU can contribute to the prediction of *i*th OTU, otherwise, there was no improvement of predicting *i*th OTU with *j*th OTU information. Therefore, the Granger causality can be tested by the ANOVA analysis to obtain a p-value. This relation between X and Y was termed as Granger causality by which implied X or Y can cause each other. Herein, the causal effects were attributed to the property of edges in the network, while OTUs would be the nodes.

### Network construction

All OTUs were filtered with two specific conditions such that OTUs with more than 80% non-zero values would be preserved, and the residuals should comply with that of at least one abundance of individual OTU reached more than 0.01% in all samples. The total number of OTUs was 98. The ADF test was applied to verify the stationarity of time series data and provide a proper lag for the next modelling process. The difference is calculated once OTUs fail in the ADF test, the results of difference will track the ADF test again. All OTUs were reserved by twice difference treatment. The time series matrix successfully inspected by the ADF test was used for the Granger test in pair. Before the operation of the Granger test, the order was been determined by VAR (R package) (Pfaff, 2008). Subsequently, the lag was transferred to the Granger test. The p-value threshold of Granger test was restricted by the following two methods. Due to the massively paired results, the links confirmed by the significance value could still cause statistical type I error, hence we introduced Bonferroni multiple-comparisons procedure and false discovery rate (FDR) to correct the threshold. Bonferroni multiple-comparisons procedure was determined by the following equation.

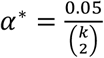

In the FDR test, all links that were selected by 0.05 significant cut-off are reordered according to the magnitude of the p-value. FDR values were calculated by the following equation.

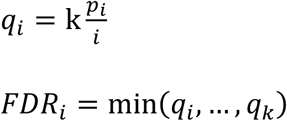

Where, *i* is the rank of the p-value in k links, which is the total links preserved by the previous threshold. The critical FDR value is normally 0.05. FDR has a great power to detect genuine positive effects, while the Bonferroni adjustment is more conservative and considers all comparisons to be statistically independent. The final file was transferred to Cytoscape software for further visualization and analysis. All analysis processes were completed with R, and several shiny apps had been built for this study (Stationary check: https://caiweiwei.shinyapps.io/stationarycheck/. Granger Causality network website: https://caiweiwei.shinyapps.io/causalnetwork/; Correlation network: https://caiweiwei.shinyapps.io/Cornetwork/; MCCN: https://caiweiwei.shinyapps.io/combinenetwork/). The specific instruction for each app is provided in the supplementary (S1).

### Network indexes

The several properties of the causal network were referenced from the literature of Anil Seth (Seth, 2005), and termed as the Causal score (*cs*), Causal density (*cd*), Net Causal flow (*ncf*), and Causal reciprocity (*c*_*recip*_). Table 1 shows equations for all corresponding properties.

As the network had been directed, outdegree and indegree represented the direction of edges within two nodes. A causal score (*cs*) was determined by the ratio of outdegree to total degrees of a specific node, reflecting the OTU influenced other OTUs rather than being influences. The causal score is defined as *cs* = the number of outdegrees divided by the number of indegrees in unweighted graphs (graphs in which all links are equivalent). If *cs* > 1, the corresponding OTU has active output, otherwise it is being passively influenced. The causal density is also termed as causal efficiency of the network, which, to some extent, represents the connectivity of the network. The net causal flow is the difference between outdegree and indegree of each node, indicating the contribution of the individual node would be either active or passive. Herein, the active state represents the species intentionally affect others, while the passive indicates it is affected by others. Although causal flow is like causal score, the former is intended to be independent of the quantity of balanced efferent and afferent connections. The causal reciprocity is the fraction of links with a directly reciprocal edge. Overall, the causal score and flow are applied to evaluate the role of each node, while the rest describes the whole network. Additionally, the supplemented indexes, including connectivity, centrality, stress centrality etc., were analyzed with the Cytoscape software tool (Feng et al., 2017).

## Data availability

All raw data were acquired from NCBI (accession number: PRJNA324303) (Jiang et al., 2018).

## ACKNOWLEDGEMENTS

This study is supported by Beijing Outstanding Young Scientist Program (BJJWZYJH01201910004016) and National Natural Science Foundation of China (NSFC, No. 51908030).

## Supporting Information

Figure S1. Description of MGCN apps family.; Figure S2. Stationary check results for 98 OTUs.; FigureS3. *ncf* values of all OTUs in MGCN.; Figure S4. Heatmap of MCN and the network of BoMCCN.; Table S1. Table of the summary of species interaction around the hub OTU56 (S5), Figure S6. MCCN of core-species; Figure S7 Node index of the specific genera from MCCN.

