## supplementary material for "Using causality and correlation analysis to decipher microbial interactions in activated sludge"

**S1 Description of MCCN app family**

The Microbial Causal Correlation Network (MCCN) apps family includes stationary check and network construction and combination app, which were coded under the framework of shiny [1]. We published the apps online with convenient accessibility for all public users (Stationary check: https://caiweiwei.shinyapps.io/stationarycheck/. Microbial Granger Causality Network (MGCN) website: <https://caiweiwei.shinyapps.io/causalnetwork/>; Microbial Correlation Network (MCN): <https://caiweiwei.shinyapps.io/Cornetwork/>; MCCN: <https://caiweiwei.shinyapps.io/combinenetwork/> ). Therefore, the shiny apps will independently work online without further requirements on local operating system or browsers, even they can be viewed by the smartphone. As the public website has been hosted by the shinyapps.io server, where users could access the features of the apps through the internet. We ensure that all apps will keep running online for more than 2 years from the publication date, which is complied with the Bioinformatics policy. The technical support will be sustainably derived from us, the error feedback will be useful for improving the quality of our website apps. There is no request on registration, all functions are freely available.

**Apps features**

As the longitudinal data was sensitive to some parameters, thereby we provided a set of apps that were described in the following content for interactive analysis. The main app is MGCN which is responsible for creating a set of MGCN files. The Stationarycheck is a satellite app for preselecting nonstationary data. The interface of main app are shown as figure 1.

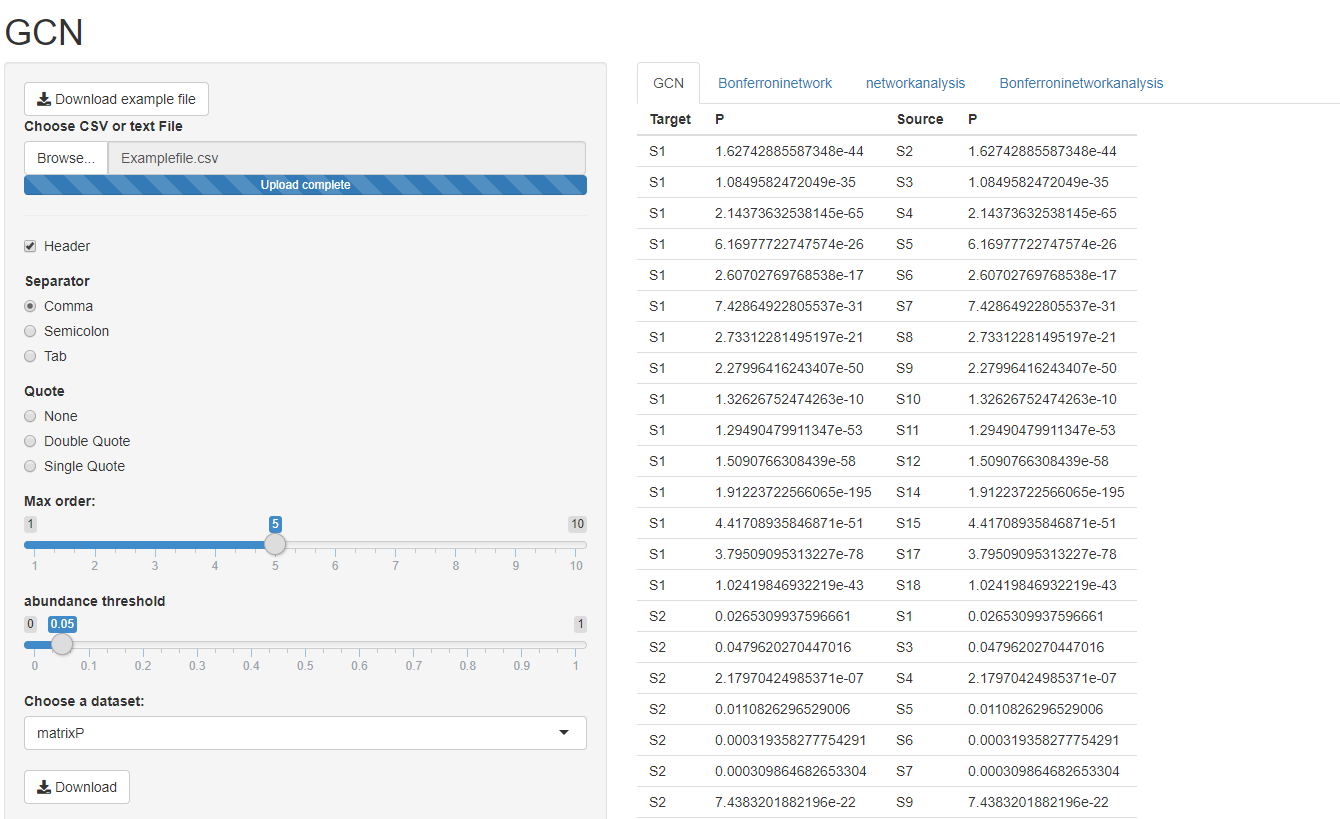

Figure1 GCN app interface

**Stationary check**

The stationary is the precondition for using the Granger test, whereas the Augmented Dickey–Fuller test (ADF test) was employed to verify if the data is satisfied with the requirement of stationary of being irrelevant to time. Herein, the difference method will be applied once if the original data do not comply with stationary. The stationary check will provide a data matrix to show the repeating number of differences before the data are subjected to the stationary. It will be important to decide which unstationary data in limited difference times should be excluded from the following analysis according to the check results.

**MGCN construction**

The main GCN construction app was composed of several modules, which include basic causal network construction, Bonferroni-correction and network analysis. The schemes of basic causal network construction were performed by the following steps details have been provided in figure 2.

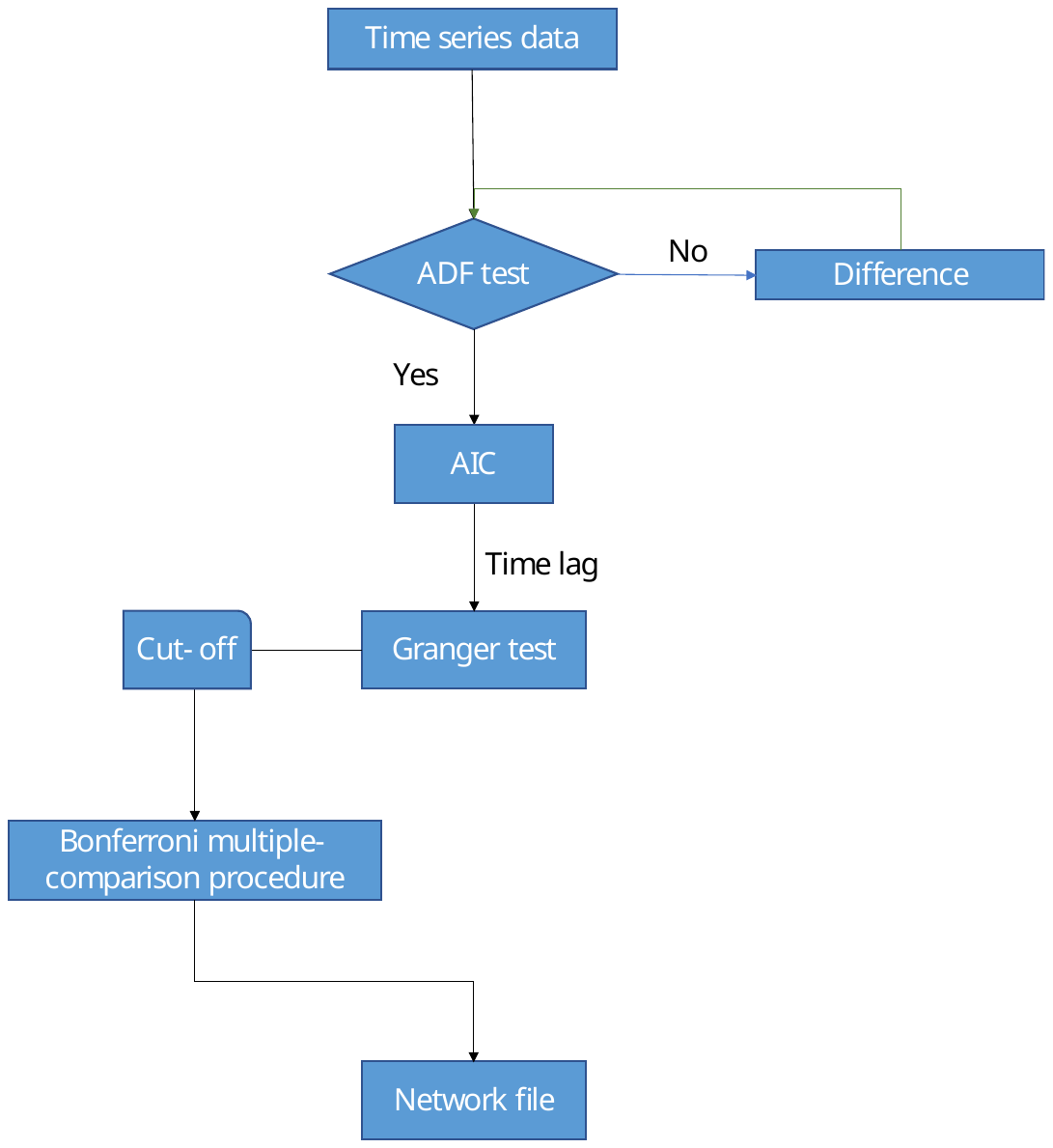

Figure 2 the pipeline of GCN app

(1) The uploaded will be filtered with the ADF test, i.e., the difference will be applied until it is stationary.

(2) Akaike information criterion (AIC) can be used to optimize the lag order for constructing the Granger test. The maximum order, representing the upper limitation of lag order decided by VAR R package, could be chosen by adjusting the “max order” function (as shown in fig 1).

(3) The basic matrix will be calculated with stationary data and optimized lag order for each paired sample. The matrix of *P* values will be filtered with a user defined cut-off value by “abundance threshold” function (as shown in fig 1).

(4) The Bonferroni-corrected threshold was developed from the cut-off value as a new standard for preserving the effective *P* matrix. As the Bonferroni-correction is extremely strict for screening the found by chance, the corrected network is a conservative one.

(5) The network analysis is founded on computing the links in a directed graph, including outdegree, indegree, neighbours, whereas to derive *cs*, *ncf*, *c_recip_* etc., which were defined in the supplementary material.

**MCN construction**

The app can be employed to construct the correlation-based network from matrix data. There will be multiple choices on the basic methods for network construction, in terms of Pearson, Spearman and Kendal. The output files include a file of MCN and a file of BoMCN, the threshold of BoMCN will be accordant with MCN.

**MCCN construction**

The final MCCN construction app can find commons from MGCN and MCN files, which will be the fundamental elements for constructing the MCCN. Two separate files separately belonged to MGCN and MCN should be uploaded, a MCCN file can be downloaded.

**S2 Figure of stationary check results for 98 OTUs**

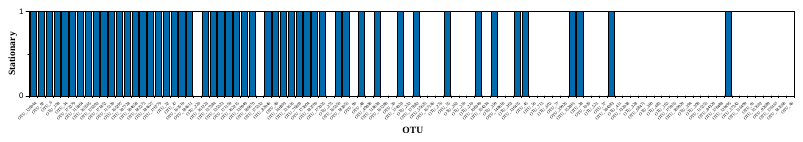

**S3 Figure of the ncf values of all OTUs in MGCN**

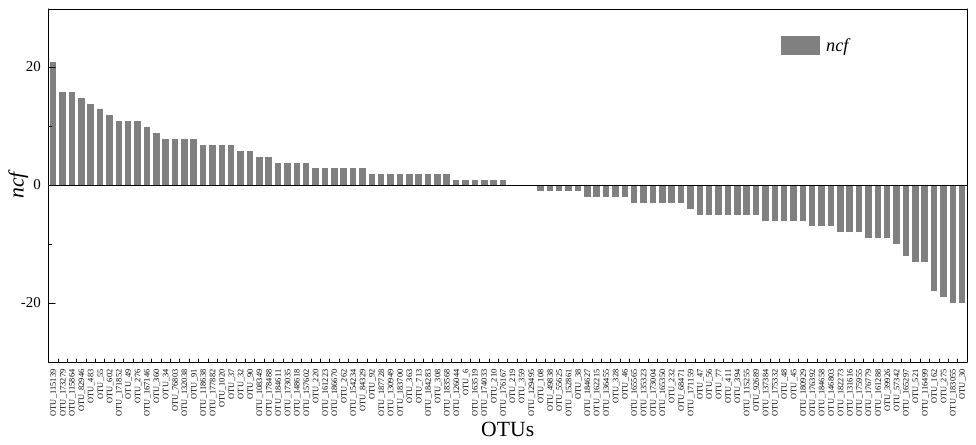

**S4 Figure of the heatmap of MCN and the network of BoMCN**

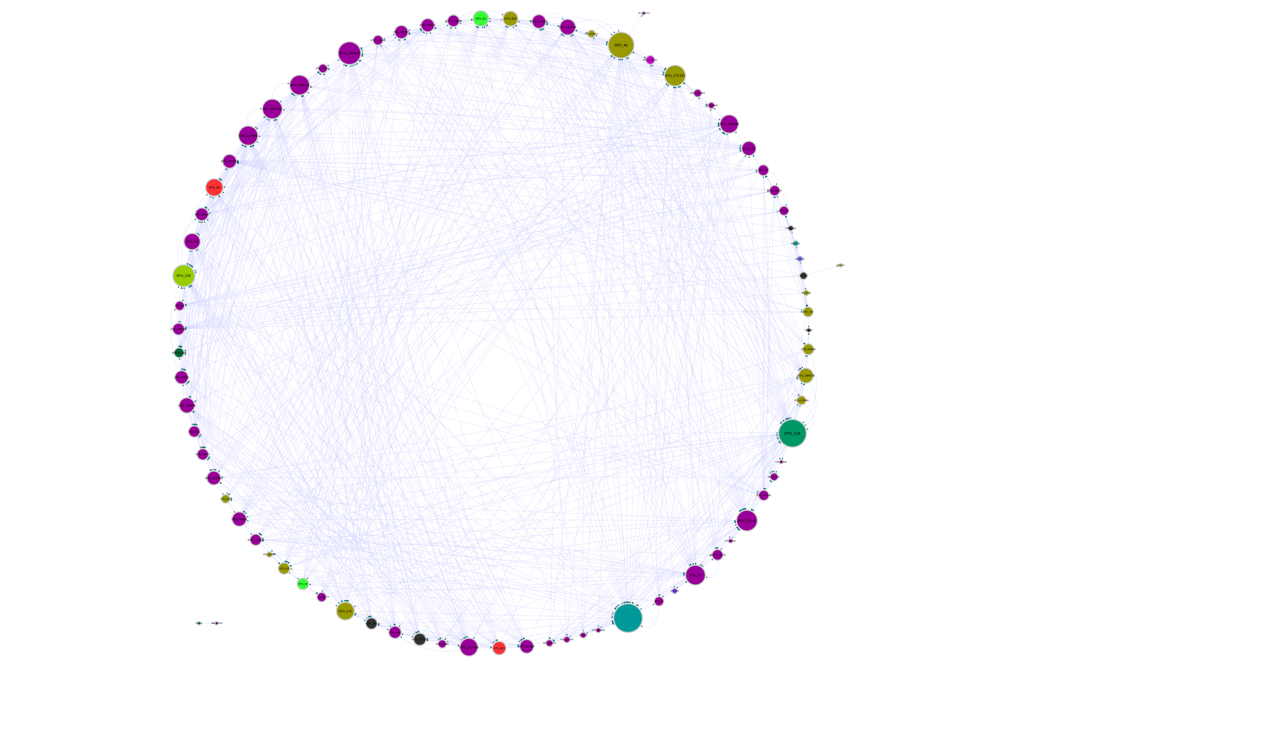

| Phylum | to | correlation | Granger casality | from | Interaction |
| --- | --- | --- | --- | --- | --- |
| Proteobacteria | OTU56 | - | unidirectional | OTU76083 | amensalism |
|  | OTU56 | - | unidirectional | OTU118638 | amensalism |
|  | OTU56 | - | unidirectional | OTU48 | amensalism |
|  | OTU56 | - | unidirectional | OTU171159 | amensalism |
|  | OTU56 | - | unidirectional | OTU161223 | amensalism |
|  | OTU56 | - | unidirectional | OTU165297 | amensalism |
|  | OTU56 | - | unidirectional | OTU176392 | amensalism |
|  | OTU56 | + | unidirectional | OTU132038 | commensalism |
|  | OTU411 | + | unidirectional | OTU56 | commensalism |
|  | OTU56 | + | unidirectional | OTU84329 | commensalism |
|  | OTU68471 | + | unidirectional | OTU56 | commensalism |
|  | OTU56 | + | unidirectional | OTU174033 | commensalism |
|  | OTU56 | + | unidirectional | OTU483 | commensalism |
|  | OTU56 | - | bidirectional | OTU173279 | competition(antagonism) |
|  | OTU56 | - | bidirectional | OTU165565 | competition(antagonism) |
|  | OTU56 | - | bidirectional | OTU131616 | competition(antagonism) |
|  | OTU56 | - | bidirectional | OTU146803 | competition(antagonism) |
|  | OTU56 | - | bidirectional | OTU135323 | competition(antagonism) |
|  | OTU56 | - | bidirectional | OTU32 | competition(antagonism) |
|  | OTU56 | - | bidirectional | OTU115139 | competition(antagonism) |
|  | OTU56 | + | bidirectional | OTU116499 | mutualism/synergism |
|  | OTU56 | + | bidirectional | OTU92689 | mutualism/synergism |
|  | OTU56 | + | bidirectional | OTU176167 | mutualism/synergism |
| *Bacteroidetes* | OTU56 | + | unidirectional | OTU275 | commensalism |
|  | OTU276 | + | unidirectional | OTU56 | commensalism |
|  | OTU56 | + | unidirectional | OTU46 | commensalism |
|  | OTU56 | - | unidirectional | OTU82946 | amensalism |
|  | OTU56 | - | unidirectional | OTU184658 | amensalism |
|  | OTU56 | - | unidirectional | OTU171852 | amensalism |
|  | OTU56 | - | unidirectional | OTU115864 | amensalism |
|  | OTU175332 | - | unidirectional | OTU56 | amensalism |
| Chlorobi | OTU56 | + | unidirectional | OTU359 | commensalism |
| Unclassified | OTU56 | - | unidirectional | OTU184611 | amensalism |
| Actinobacteria | OTU56 | - | unidirectional | OTU55 | amensalism |
| Chloroflexi | OTU56 | + | unidirectional | OTU602 | commensalism |
| Firmicutes | OTU394 | + | unidirectional | OTU56 | commensalism |
|  | OTU175955 | + | unidirectional | OTU56 | commensalism |

**S5 Table of the summary of species interaction around the hub OTU56**

**S6 MCCN of hub-species**

Each color represents individual phylum according to the taxonomy of microorganisms. The size of the node and node label is linearly proportionate to the edge number of each node from 0 to 50. The arrows represent the direction of Granger causality. The pink color of the link represents a positive association and grey link is the negative connection. The size of the link is linearly proportionate to the correlation absolute value from 0 to 1. All bars represent the composition of interaction types between OTU56 with other taxa.

**S7 Figure of nodes index of specific genus from MCCN**

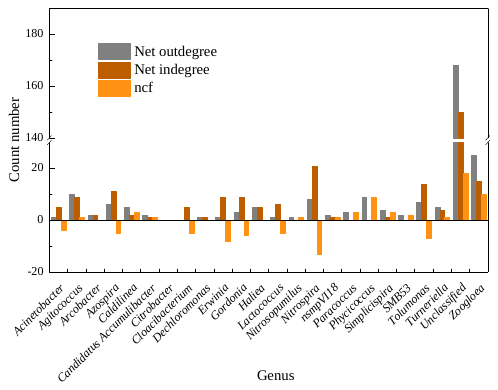

1.	Chang W, Cheng J, Allaire JJ, Xie Y, McPherson J. Package ‘shiny’. *See Httpciteseerx Ist Psu Eduviewdocdownload* 2015.
